# Dried Blood Spot Lipidomics in Diabetes

**DOI:** 10.1101/2025.05.28.656742

**Authors:** Jayden Lee Roberts, Monique J. Ryan, Luke Whiley, Nicola Gray, Melvin Gay, Elaine Holmes, Jeremy K. Nicholson, Julien Wist, Nathan G. Lawler

## Abstract

Comprehensive lipidomic profiling in diabetes has identified disease-associated lipid signatures that may support more personalised monitoring beyond routine glycaemic control. Dried blood spot (DBS) microsampling offers a minimally invasive method for collecting and storing small blood volumes (<50 µL), enabling decentralised and longitudinal sampling. However, its application in lipidomics for cardiometabolic phenotyping remains underexplored. In this study, 34 participants (17 diabetics, 17 non-diabetic controls) self-collected DBS samples using advanced microsampling devices (10 µL, Capitainer®B). Lipid extracts were analysed using a validated liquid chromatography–trapped ion mobility–time-of-flight mass spectrometry method optimised for semi-quantitative DBS lipidomics. Multivariate analysis revealed significant differences in lipidomic profiles (n = 432 lipids) between diabetic and non-diabetic individuals (R²Y = 0.82, Q²Y = 0.50, p = 0.05). Key discriminatory lipids (all VIP > 1.44), including HexCer(18:1;O2/24:0), HexCer(18:1;O2/22:0), LPC(16:0), PC(O-34:3), TG(18:0_18:1_20:3), and TG(18:1_18:1_22:6), were consistent with matched venous and capillary plasma samples. Lipid class and structural differences, including elevated long-chain triacylglycerols and reduced lysophosphatidylcholines reflected known dyslipidaemia in diabetes and demonstrate the capacity of DBS to capture biologically meaningful lipid perturbations. Participant-reported outcomes from Problem Areas In Diabetes (PAID-20), Patient Activation Measure (PAM-13), and microsampling perception questionnaires highlighted psychosocial and behavioural insights, and indicated strong support for DBS sampling over venepuncture, citing ease of use and reduced burden. These findings establish DBS lipidomics as a feasible and informative approach for stratifying cardiometabolic disease, supporting broader implementation in remote, personalised monitoring frameworks (Figure 1).

**Figure 1.**
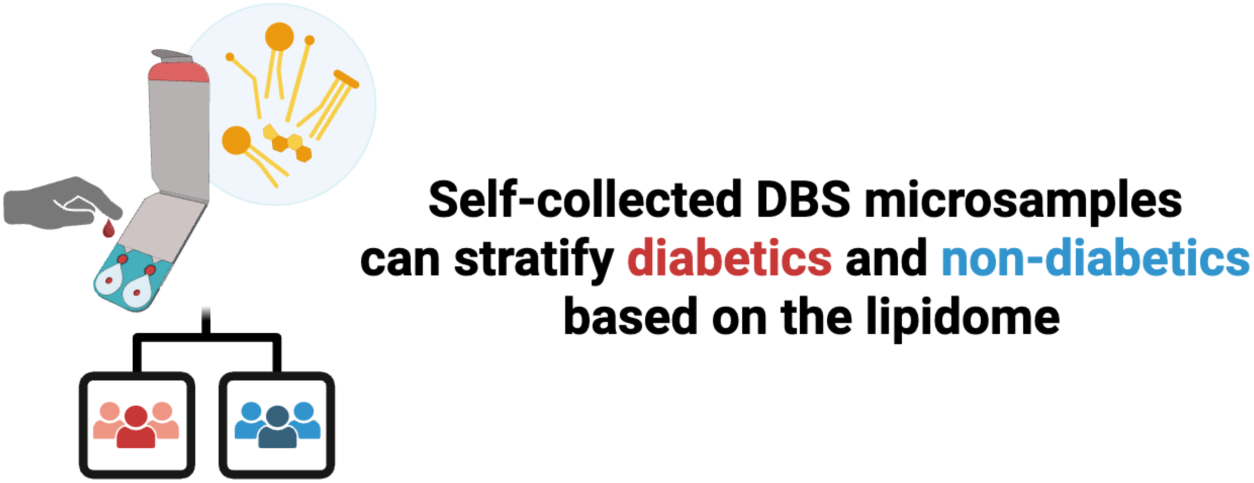
**Graphical abstract – Chapter 5**.

## INTRODUCTION

Diabetes mellitus is a growing global health challenge with an estimated prevalence of 830 million individuals affected worldwide^1^. Disruptions in lipid composition and metabolism^2^, are central to the pathophysiology of diabetes, contributing to complications including insulin resistance, diabetic kidney disease, retinopathy, neuropathy, metabolic dysfunction, and disease progression^3^. For instance, alterations in glycerophospholipids (e.g., lysophosphatidylcholines)^4–7^, sphingolipids (e.g., ceramides)^8–11^, and glycerolipids (e.g., triacylglycerols)^12–14^, have been shown to contribute to insulin resistance and cardiovascular risk. Advances in lipidomics now offer new opportunities to elucidate the complex interplay between lipid perturbations and metabolic dysfunction, with potential to improve biomarker discovery and personalise diabetes management. Patient-centric sampling, combined with lipidomics enables minimally invasive profiling while offering mechanistic insights into lipid alterations linked to disease onset, progression, and complications to support early detection, risk stratification, and personalised intervention in diabetes.

Microsampling techniques, such as finger-stick capillary blood collection (<50 µL), offer a practical alternative to traditional venous blood sampling, addressing the need for accessible and reproducible health assessments in management of chronic diseases such as diabetes. These methods support more frequent, patient-friendly lipid profiling, aligning with the principles of predictive, preventive, personalised, and participatory (P4) medicine^15^. Microsamples can be collected as whole blood collected (to derive capillary plasma and serum), or as dried blood spots (DBS)^16^. DBS have been used in neonatal screening of inborn errors of metabolism, such as phenylketonuria, since the 1960s^17^. DBS offer a minimally-invasive, cost-effective approach that circumvents challenges associated with venous sampling, including the need for clinical appointments, trained personnel, and cold-chain logistics. As patients with diabetes are already accustomed to remote health assessments, such as spot or continuous glucose monitoring (CGM), integrating DBS sampling represents a logical extension of current care models^15, 18, 19^. And by integrating DBS sampling for parallel assessment of lipids, this may provide additional insight into metabolic risk, disease progression, and therapeutic response beyond glycaemic control.

DBS-based analytical approaches such as liquid chromatography-mass spectrometry (LC-MS) can detect subtle molecular alterations, offering a powerful tool to characterise metabolic dysregulation in diabetes beyond traditional venous sampling of routine clinical markers^20^. For example, DBS have previously undergone investigation alongside CGM for measurement of clinical markers such as glycated haemoglobin (HbA1c)^21–26^ and c-peptide^27^. In addition to metabolomic profiling, DBS have also been applied in preclinical models for protein-level analysis using targeted multiplex proteomic panels^28^. Similarly, DBS microsampling could advance our understanding of lipid dysfunction in diabetes by enabling longitudinal tracking of lipidomic changes in response to lifestyle modifications, medication, or disease progression^29^. However, despite the growth of lipidomics in diabetes research, most studies continue to rely on venous sampling, limiting the real-world applicability of lipidomic profiling. Thus, leveraging DBS lipidomics alongside traditional glucose monitoring and clinical markers such as HbA1c, could shift diabetes management toward a more holistic and decentralised health model, allowing for personalised metabolic insights and earlier intervention strategies tailored to individual patients.

In the present study, we analysed and compared lipid profiles across venous plasma, capillary plasma, and whole blood DBS using a validated untargeted lipidomics platform for liquid chromatography– trapped ion mobility spectrometry–time-of-flight mass spectrometry (LC–timsTOF–MS).

## EXPERIMENTAL SECTION

### Reagents and chemicals

Optima LC-MS grade organic solvents were purchased from Thermo Fisher Scientific (Malaga, WA, Australia). LC-MS grade water was generated from a Milli-Q IQ 7000 (Merck Millipore). EquiSPLASH® LIPIDOMIX® stable isotopically labelled internal standard (SIL-ISTD) mixture was obtained from Avanti Polar Lipids (Sigma-Aldrich, North Ryde, NSW, Australia). The working solution of SIL-ISTD extraction solvent was created by diluting EquiSPLASH® 1:100 with propan-2-ol (IPA) for plasma samples and 80 % IPA in water for DBS samples, as described elsewhere^30^.

### Experimental design

The study comprised four parts: i) Participant recruitment (see Section 5.3.3: ‘Participants’), ii) Clinical visit (see Section 5.3.4: ‘Surveys’ and Section 5.3.5: ‘Biological sample collection’), iii) Biological sample analysis (see Section 5.3.6.1: ‘Biological sample preparation’ and Section 5.3.6.2: ‘High-resolution untargeted lipidomics’), and iv) Statistical analysis of biological sample data (see Section 5.3.7: ‘Statistical analysis’) (Figure 2).

**Figure 2.**
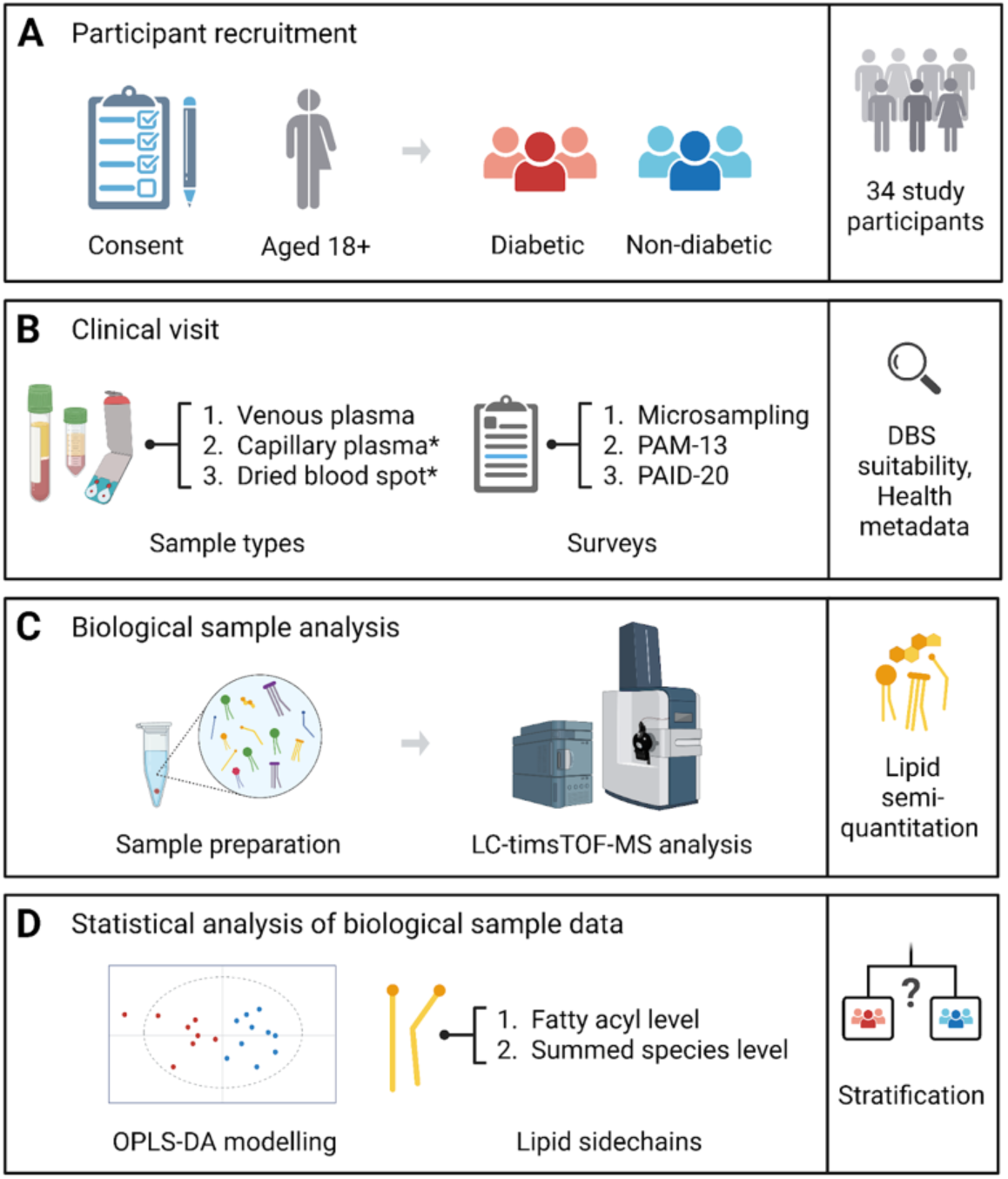
Generalised workflow of participant recruitment, sample collection, lipid profiling, and statistical analysis. (A) Enrolment criteria, with final participant number used in this study (n = 34). (B) Biological sampling included *self-collected dried blood spots (Capitainer®B), venous plasma, and capillary plasma. Surveys included perception of microsampling, PAM-13, and PAID-20. (C) Simplified untargeted lipid profiling analysis using LC-timsTOF-MS to detect 432 lipid species spanning 25 subclasses. (D) Statistical analysis comparing lipidomic profiles of diabetic (n = 17) and non-diabetic control (n = 17) participants using OPLS-DA modelling, and lipid side chain profiling. Image created using BioRender.com.

### Participants

A total of 34 participants (age range: 21 - 80 years) were recruited, comprising 17 individuals with diabetes (mean age: 48.7 ± 14.2 years) and 17 non-diabetic controls (mean age: 44.1 ± 14.2 years). Diabetes status was defined as fasting plasma glucose ≥ 7.0 mmol/L, HbA1c ≥ 6.5%, or use of antidiabetic medication. Non-diabetic controls were defined as having fasting plasma glucose < 5.6 mmol/L, HbA1c < 5.7%, or no history of diabetes. Eligibility for the study was deemed as anyone who was over 18 years of age, willing to provide venous and capillary blood samples, and complete a health questionnaire and surveys (perceptions of microsampling survey based on Van Uytfanghe et al.^31–33^, Patient Activation Measure-13 (PAM-13)^34^, and Problem Areas in Diabetes-20 (PAID-20)^35^). A written description of the study’s risks and benefits was given to participants, who provided written informed consent before the study’s commencement. This study was approved by Murdoch University’s Human Research Ethics Committee (Ethics no. 2022/119) in compliance with the Australian National Statement and the Declaration of Helsinki.

### Surveys

Survey data was collected from participants. This included a survey on perceptions of microsampling, the PAM-13, and PAID-20. Only diabetic participants completed the PAID-20, as this is a survey specific to diabetes.

#### Perception of microsampling survey

A 9-question survey assessing perception of microsampling based on the VAMS microsampler in a study by Van Uytfanghe et al. (2021)^31^ was adapted for Capitainer®B microsamples. The survey assesses aspects of microsampling including clarity of collection instructions, user-friendliness, preference for collection in a home setting compared to traditional phlebotomy, frequency participants normally perform finger-pricks, and their perceived pain of the finger-prick. Participants completed the perceptions of microsampling survey following self-collection of their microsample.

#### PAM-13

The Patient Activation Measure-13 is a validated survey that assesses a patient’s knowledge, skills, and confidence in managing care of their health^34^. It comprises 13 items, scored using Likert scales to classify individuals into four levels of increasing activation: ‘disengaged and overwhelmed’, ‘becoming aware’, ‘taking action’, and ‘maintaining behaviors’. Higher PAM-13 scores indicate greater patient activation, which is linked to better health outcomes, improved adherence to treatment, and proactive health management. The PAM-13 has previously been utilised in the context of diabetes^36–39^.

#### PAID-20

The Problem Areas in Diabetes-20 is commonly used in diabetes research and clinical practice to identify patients struggling with the emotional aspects of diabetes and to provide targeted psychological and behavioural support^35, 40^. It comprises 20 questions designed to assess diabetes-related emotional distress. It measures the psychosocial burden of living with diabetes, including frustration, guilt, fear of complications, and difficulties with self-management. Each item is scored on a 5-point Likert scale (0 = Not a problem, 4 = Serious problem), with total scores ranging from 0 to 100 (after multiplying the sum by 1.25). Higher scores (> 40) indicate clinically significant diabetes distress that may require intervention.

### Biological Sample Collection

Participants attended a clinic following a 12-hour overnight fast for the collection of a venous sample and capillary blood samples (capillary plasma and DBS). A trained phlebotomist performed venous blood collection using a 23-gauge needle into a Becton Dickinson (BD) vacutainer tube taken from the antecubital fossa (1 × 10 mL of lithium heparin plasma tube). Capillary blood microsampling utilised commercially available Capitainer®B (Capitainer, Solna, Sweden) advanced microsampling devices for collection of DBS samples from the fourth digit (finger) by the study participants using a BD microtainer contact-activated lancet (Becton Dickinson, New Jersey, United States). Each device collected duplicate 10 µL capillary blood DBS microsamples. The remaining capillary blood droplets were collected into a MiniCollect® 0.5 mL lithium heparin plasma tube (Greiner Bio-One, Austria). Prior to microsample collection, participants were provided with an instructional pamphlet^41^ and instructional video^42^ from the manufacturer. Venous and capillary samples were collected within 20 minutes. Venous and capillary blood tubes were centrifuged at 4 °C for 10 min at 1300 x g. The resulting plasma layers were aliquoted into cryo-vials and stored at −80 °C until lipidomic analysis. DBS capillary samples were dried at room temperature for 4 hr before being stored at −80°C.

### Lipidomic Analysis

#### Biological sample preparation

Sample preparation utilised pre-validated monophasic solvent extraction methodologies described for plasma^43^ and whole-spot DBS microsamples^30^. Monophasic extraction solvents contained SIL-ISTD (EquiSPLASH® LIPIDOMIX®) at a dilution of 1 µg/mL. 20 µL plasma samples were extracted with 180 µL of IPA and 10 µL DBS were extracted with 150 µL of 80 % IPA in water. All samples underwent a 10 min vortex mix, with DBS samples undergoing an additional 5 min sonication after vortexing. Following this, samples underwent a 20 min protein precipitation at −20 °C, and 10 min centrifugation at 14000 x *g* (4 °C). The supernatant was then collected and split across two 350 μL 96-well plates (Eppendorf, Macquarie Park, NSW, Australia) for LC-timsTOF-MS analysis in two ionisation modes: positive and negative heated electrospray ionisation modes. Pooled quality control (PQC) samples were created by combining equal volumes of sample extracts.

#### High-resolution untargeted lipidomics

A high-resolution untargeted lipid profiling approach for plasma and DBS microsamples was employed following a previously published LC-timsTOF-MS method^30^ to detect lipid species from 25 lipid subclasses. Briefly, untargeted lipid profiling was performed using a Waters I-Class UPLC system (Waters Corporation, MA, USA) coupled to a Bruker timsTOF Pro (Bruker, MA, USA) equipped with a vacuum insulated probe-HESI source. The analytical parameters have been detailed in the supporting information (See Appendix D.1).

#### Data processing

LC-timsTOF-MS raw data files were pre-processed using commercially available software Metaboscape 2024b (Bruker, MA, USA) to perform peak picking, rule-based *in silico* lipid annotation using lipid-match algorithms (precursor *m/z*, isotopic pattern, characteristic fragments in MS/MS spectra) and measured collisional cross section (CCS) values as an additional qualifier. Data post-processing was performed using R v4.4.1 and RStudio v1.4.1^44^. This included lipid semi-quantitation which was performed by calculating the relative concentration of each lipid analyte based on its ratio to a surrogate SIL-ISTD within the same subclass per Roberts et al. (2025)^30^.

#### Statistical analysis Survey data

Independent t-tests were performed using R v4.4.1 and RStudio v1.4.1^44^ to assess for significant differences in survey responses between diabetic participants and non-diabetic controls for the perceptions of microsampling survey and the PAM-13. Stacked bar plots were generated using *ggplot2*^45^ and *likert*^46^ to visualise spread of survey responses for each question, respectively, perceptions of microsampling, PAM-13, and PAID-20.

#### Biological sample data

Prior to statistical analysis, samples or lipids with over 50 % missing or zero values across the samples within the total dataset were filtered out. The remaining missing values were then imputed (min/2), and a 30 % coefficient of variation (CV) filter was applied on replicate PQC samples. Orthogonal projections to latent structures discriminant analysis (OPLS-DA) modelling was performed to derive variable importance in projection (VIP) scores using the *mva.plots* package (V 0.0.8)^47^. Lipids with VIP scores exceeding 1.44 were considered relevant for class separation, based on the minimum observed value among discriminant features consistently identified across all OPLS-DA models. Lipid side chain plots were constructed in R, using *ggplot2*, to visualise differences between lipidomic profiles of diabetics and non-diabetic controls, similar to the *lipidomeR*^48^ approach by comparing acyl chain length and degree of lipid species unsaturation. Here, lipids were categorised based on acyl chain composition using shorthand notation (total number of carbons : degree of unsaturation). This was performed at two levels: fatty acyl level, and summed species level.

## RESULTS

The participant demographics, including age, sex, weight, height, BMI are shown in Table 1. Responses between diabetic participants and non-diabetic controls for the perceptions of microsampling survey and the PAM-13 are also reported in Table 1, and PAID-20 results for diabetics reported in Table S.1. The spread of Likert responses between diabetic participants and non-diabetic controls for the perceptions of microsampling survey are shown in Figure 3, and responses for the PAM-13 are presented in Figure 4. Likert responses from diabetic participants for the PAID-20 survey are presented in Figure 5.

**Figure 3.**
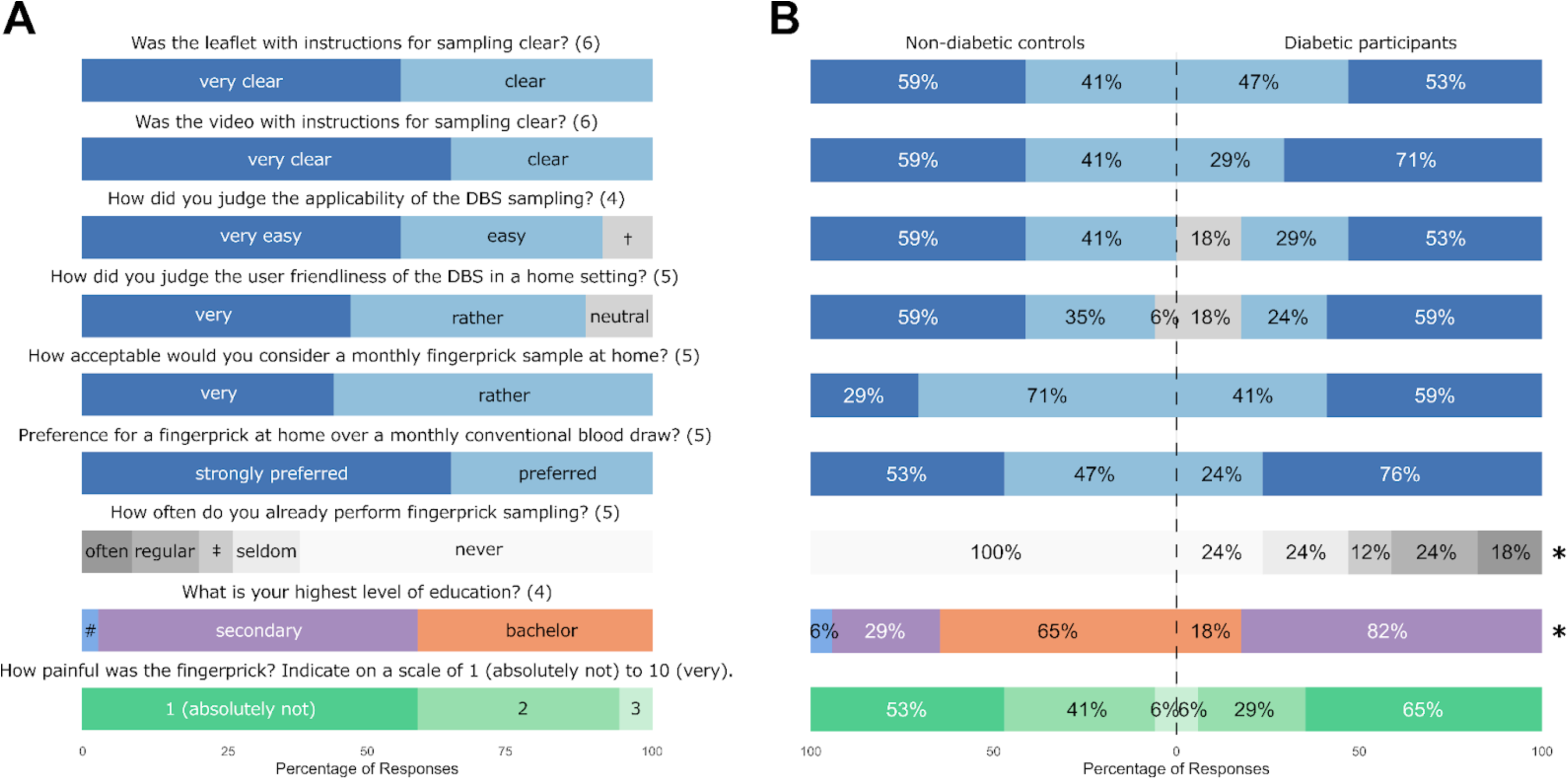
Perceptions of microsampling survey Likert scale responses. (A) all study participants, and (B) study participants by group (non-diabetic controls vs. diabetic participants). Numbers in brackets represent the total number of Likert scale options available for the specific question. Nb. not all responses were selected by study participants. †neither easy, nor difficult. ‡every now and then. #primary. *Survey questions with significant differences in response between diabetics and non-diabetic controls.

**Figure 4.**
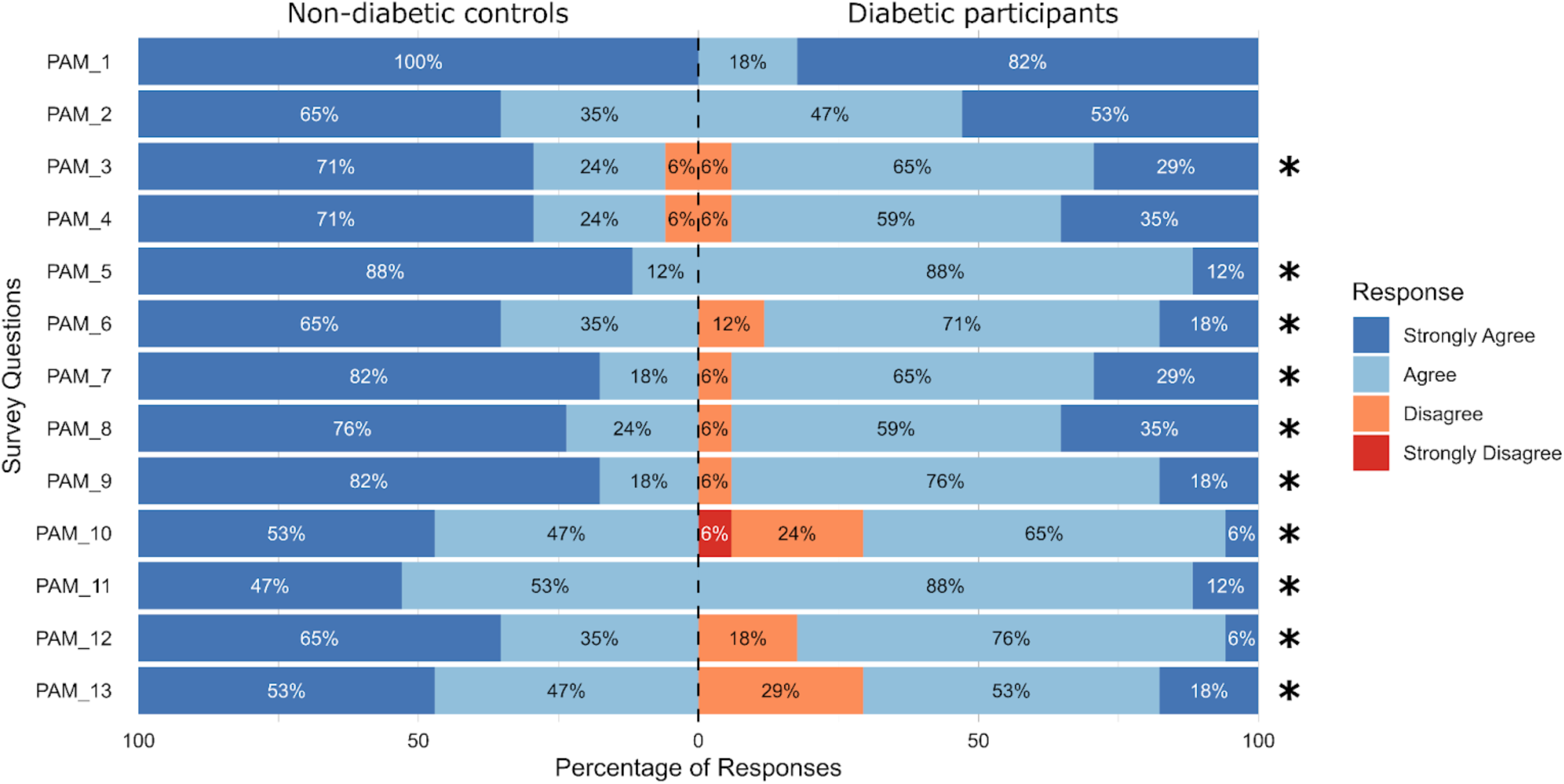
Patient Activation Measure (PAM-13) Likert scale responses. “PAM_1” refers to question 1 of the survey, as seen in Table 1. *Questions with significant differences between diabetics and non-diabetic controls.

**Figure 5.**
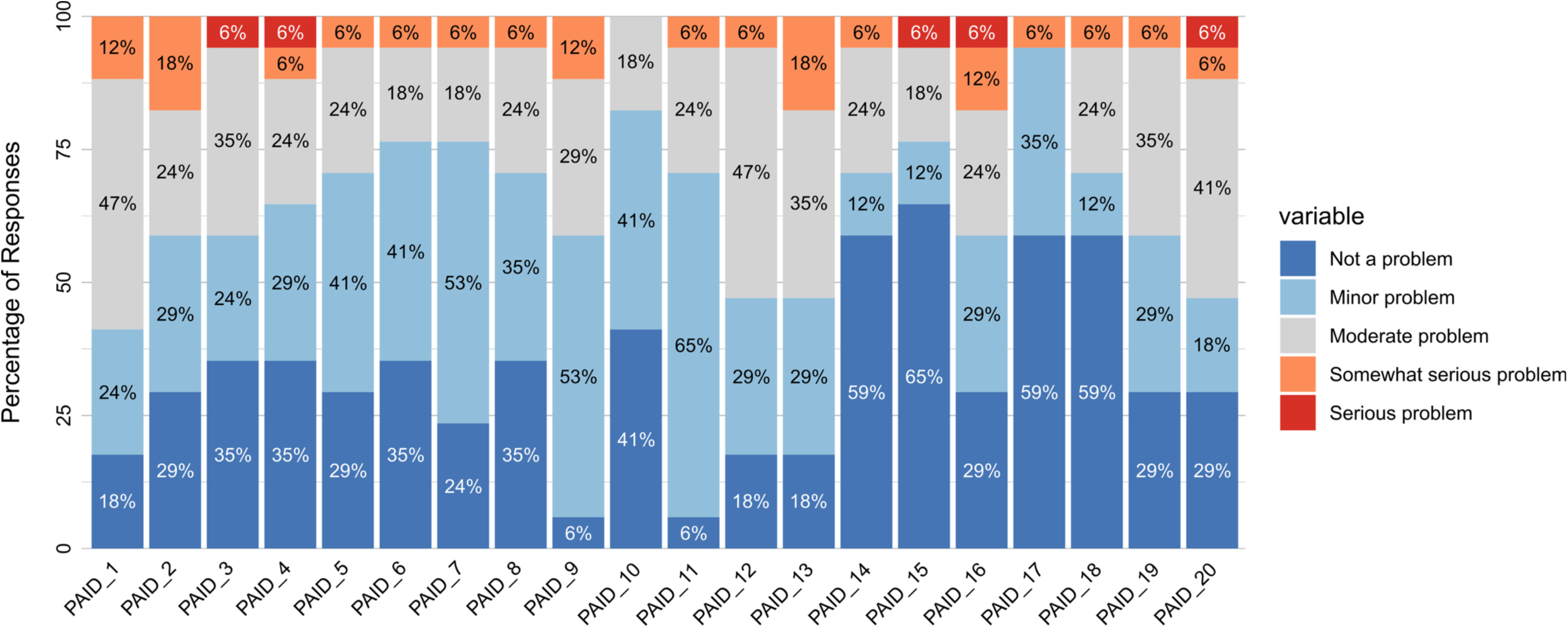
Problem Areas in Diabetes (PAID-20) Likert scale responses for diabetic participants. “PAID_1” refers to question 1 of the survey. A descriptive list of the abbreviated PAID-20 survey items are reported in Table S.1.

**Table 1.** Participant demographics.

|  | Non-diabetic (n = 17) | Diabetic (n = 17) | Sig. |
| --- | --- | --- | --- |
| **Sex - Female (%)** | n = 9 (53%) | n = 9 (53%) | p = 1 |
| **Age (years)** | 44.8 [±14.2] | 48.2 [±14.1] | p = 0.3699 |
| **Weight (kg)** | 68.4 [±11.3] | 76.1 [±14.9] | p = 0.0996 |
| **Height (cm)** | 165.8 [±11.5] | 164.4 [±9.8] | p = 0.7080 |
| **BMI** | 25 [±4.1] | 28.3 [±6.1] | p = 0.0762 |
| **Fasting glucose** | 5.2 [±0.7] | 8.8 [±1.7] | **p < 0.0001** |
| **HbA1C** | - | 8.44 [±1.55] |  |
| Good glycaemic control (HbA1C <7%) | - | n = 2 |  |
| Poor glycaemic control (HbA1C ≥7%) | - | n = 15 |  |
| **Perceptions of microsampling** |  |  |  |
| 1. Leaflet instructions clear | 1.41 [±0.51] | 1.47 [±0.51] | p = 0.7393 |
| 2. Video instructions clear | 1.41 [±0.51] | 1.29 [±0.47] | p = 0.4880 |
| 3. Applicability of DBS sampling | 1.41 [±0.51] | 1.65 [±0.79] | p = 0.3075 |
| 4. User-friendliness | 1.71 [±0.59] | 1.59 [±0.8] | p = 0.6271 |
| 5. Acceptability of monthly finger-prick | 1.71 [±0.47] | 1.41 [±0.51] | p = 0.0890 |
| 6. Preference for finger-prick over blood draw | 1.47 [±0.51] | 1.24 [±0.44] | p = 0.1605 |
| 7. Finger-prick frequency | 1 [±0] | 2.88 [±1.5] | **p < 0.0001** |
| 8. Education | 2.59 [±0.62] | 2.18 [±0.39] | **p = 0.0270** |
| 9. Pain | 1.53 [±0.62] | 1.41 [±0.62] | p = 0.5847 |
| **PAM-13** |  |  |  |
| 1. Responsible for taking care of my health | 4 [±0] | 3.82 [±0.39] | p = 0.0733 |
| 2. Taking active role in health | 3.65 [±0.49] | 3.53 [±0.51] | p = 0.5008 |
| 3. Confident I can reduce problems | 3.65 [±0.61] | 3.24 [±0.56] | **p = 0.0483** |
| 4. Know what my prescribed medications do | 3.65 [±0.61] | 3.29 [±0.59] | p = 0.0945 |
| 5. See doctor vs take care of problem myself | 3.88 [±0.33] | 3.12 [±0.33] | **p < 0.0001** |
| 6. Tell concerns to doctor when they do not ask | 3.65 [±0.49] | 3.06 [±0.56] | **p = 0.0026** |
| 7. Follow through on medical treatments at home | 3.82 [±0.39] | 3.24 [±0.56] | **p = 0.0013** |
| 8. Understand my health problems and their cause | 3.76 [±0.44] | 3.29 [±0.59] | **p = 0.0125** |
| 9. Know treatments available for health problems | 3.82 [±0.39] | 3.12 [±0.49] | **p = 0.0001** |
| 10. Can maintain lifestyle changes (diet, exercise) | 3.53 [±0.51] | 2.71 [±0.69] | **p = 0.0004** |
| 11. Know how to prevent health problems | 3.47 [±0.51] | 3.12 [±0.33] | **p = 0.0236** |
| 12. Figure out solutions for new health problems | 3.65 [±0.49] | 2.88 [±0.49] | **p = 0.0001** |
| 13. Maintain lifestyle changes during stress | 3.53 [±0.51] | 2.88 [±0.7] | **p = 0.0042** |
| **PAM-13 subscales** |  |  |  |
| A belief that role is important (Items 1+2) | 7.65 [±0.49] | 7.35 [±0.79] | p = 0.2004 |
| Having confidence and knowledge (Items 3-8) | 22.41 [±1.28] | 19.24 [±2.28] | **p < 0.0001** |
| Taking action (Items 9-11) | 10.82 [±0.88] | 8.94 [±0.83] | **p < 0.0001** |
| Staying on course under stress (Items 12+13) | 7.18 [±0.95] | 5.76 [±1.09] | **p = 0.0003** |
| **Patient activation level n(%)** |  |  |  |
| 1 (0-47) | 0 (0%) | 0 (0%) |  |
| 2 (47.1-55.1) | 0 (0%) | 4 (24%) |  |
| 3 (55.2-72.4) | 0 (0%) | 8 (47%) |  |
| 4 (72.5-100) | 17 (100%) | 5 (29%) |  |

*Note.* Statistical testing involved Chi-square for sex assessment. Significant p-values derived from independent t-tests are shown in bold.

### Survey analysis

For the perceptions of microsampling survey, on average, participants indicated that microsampling instructions were “very clear” (both pamphlet and video). Applicability in home settings was reported “very easy” and user-friendliness was rated as “very”. Preference over a traditional phlebotomy blood draw was reported as “strongly preferred”, and pain from the finger-prick was reported as <2 on a 10-point scale. Significant differences were identified between diabetic participants and non-diabetic controls for question 7 “How often do you already perform finger-prick sampling?” t(32) = 5.19, p < 0.0001 and question 8 “What is your highest level of education?” t(32) = −2.32, p = 0.0270.

Significant differences were identified between PAM-13 survey responses of participants with diabetes and non-diabetic controls. These mainly concerned survey items related to domains of having confidence and knowledge (summed items: 3-8) (t(32) = −5.01, p < 0.0001), taking action (summed items: 9, 11) (t(32) = −6.42, p < 0.0001), and staying on course under stress (summed items: 12, 13) (t(32) = −4.02, p < 0.0003). Non-diabetic controls reported higher activation levels for these parameters.

Parameters of the PAID-20 survey where diabetic participants identified serious problems included item 3: *scared of living with diabetes*, 4: *uncomfortable in social situations* (e.g., telling people what you eat), 15: *unsatisfied with physician*, 16: *diabetes taking up too much mental and physical energy*, and 20: *feeling “burned out” by diabetes management*. These parameters largely pertain to the diabetes-related emotional distress domain.

### Lipidomic analysis

One sample was removed from the DBS data set prior to any statistical analysis due to a failed quality control check, where less than the required 10 μL of blood was collected. In total, 23/25 lipid subclasses and 432 lipid species were semi-quantified in plasma and DBS samples using a pre-validated LC-timsTOF-MS method^30^.

OPLS-DA modelling was performed using all annotated lipids (n = 432) (Figure 6). Significant differences were identified between DBS samples of diabetic individuals and non-diabetic controls (R²Y = 0.82, Q²Y = 0.50, p = 0.05), indicating that a substantial proportion of class variation was explained by the model. This observation was consistent with matched traditional venous plasma samples (R²Y = 0.77, Q²Y = 0.53, p = 0.05) and capillary plasma samples (R²Y = 0.67, Q²Y = 0.44, p = 0.05). The OPLS-DA loadings plot of the DBS samples revealed the lipids driving this separation (see Figure S.1 and Figure S.2), and top VIP scores indicated HexCer(18:1;O2/24:0), HexCer(18:1;O2/22:0), LPC(16:0), PC(O-34:3), TG(18:0_18:1_20:3), and TG(18:1_18:1_22:6) were consistent in stratifying diabetics and non-diabetic controls for all three sample types (all VIP > 1.44) (Figure 6).

**Figure 6.**
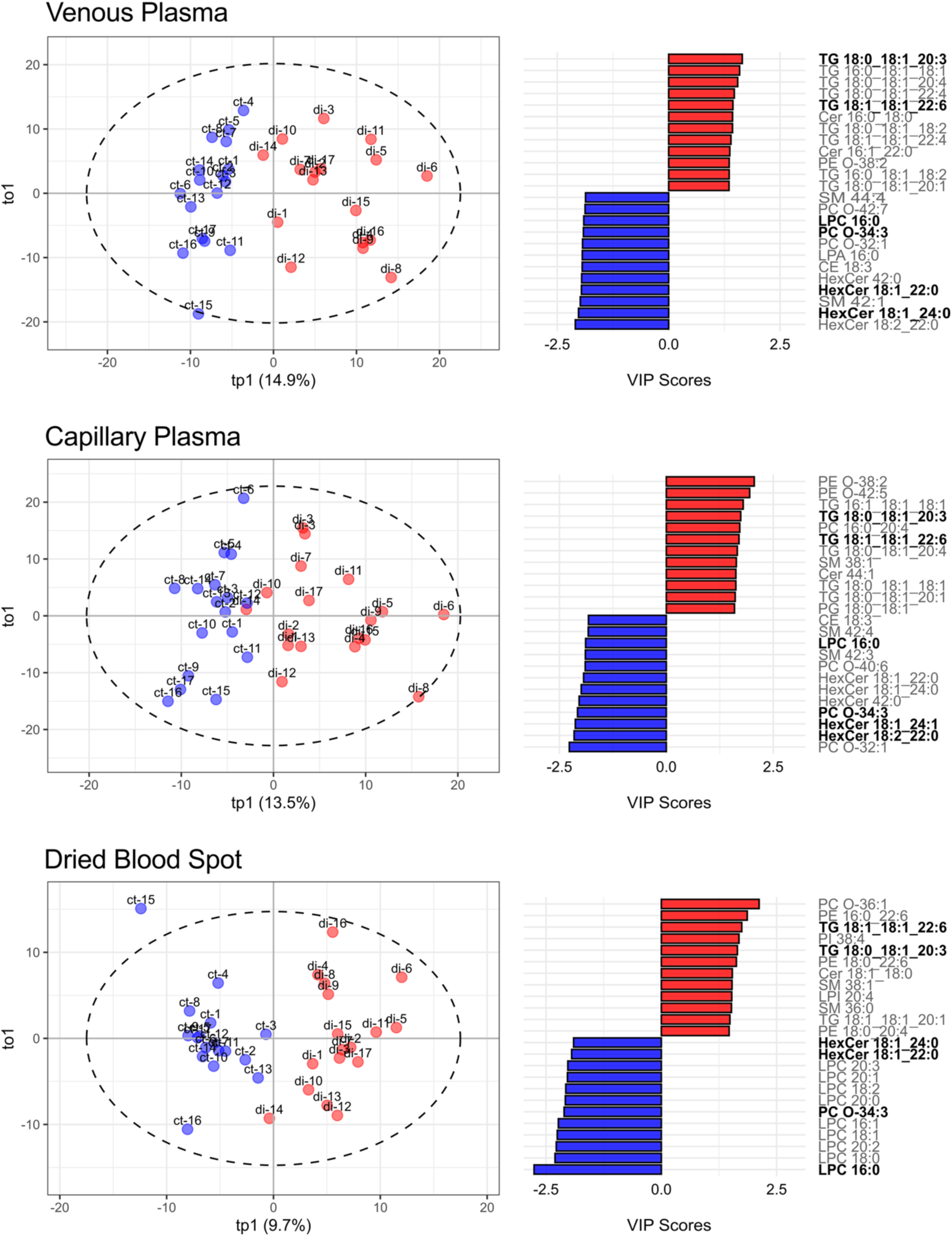
OPLS-DA scores and top VIP-ranked lipids discriminating diabetic and control samples across biofluids. Scores plot of supervised OPLS-DA modelling for non-diabetic controls (n = 17) and diabetics (n = 17) by biofluid (venous plasma, capillary plasma, dried blood spots). Bar plots correspond to the top 12 loading variables of the DBS OPLS-DA model (ranked by VIP score), colours represent diabetic participants (red) vs non-diabetic controls (blue). Lipids in bold indicate those that were in the top VIP scores across all three biological samples (venous plasma, capillary plasma, and dried blood spot).

The VIP scores derived from the DBS OPLS-DA model, were used to visualise the lipids contributing to group separation between diabetic participants and non-diabetic controls. To investigate how individual acyl chain length and degree of unsaturation influence diabetes, DBS lipids were examined at the fatty acyl level (Figure 7). Glycerolipids such as triacylglycerol (TG) species demonstrated notable enrichment in diabetic participants, particularly for lipid species with longer and more unsaturated side chains: for example, TG species containing acyl chains C22:6 and C20:3. Notably, LPC species uniformly have higher VIP scores in non-diabetic controls. Lipids with shorter, predominantly saturated and monounsaturated fatty acyl side chains displayed stronger associations with non-diabetic controls across multiple subclasses. For example, C14:0 – myristic acid, C16:0 – palmitic acid, C16:1 – palmitoleic acid, C18:0 – stearic acid, and C18:1 – oleic acid. A similar trend was seen for polyunsaturated C18 acyl chains including C18:2 – linoleic acid, and C18:3 – alpha-linoleic acid.

**Figure 7.**
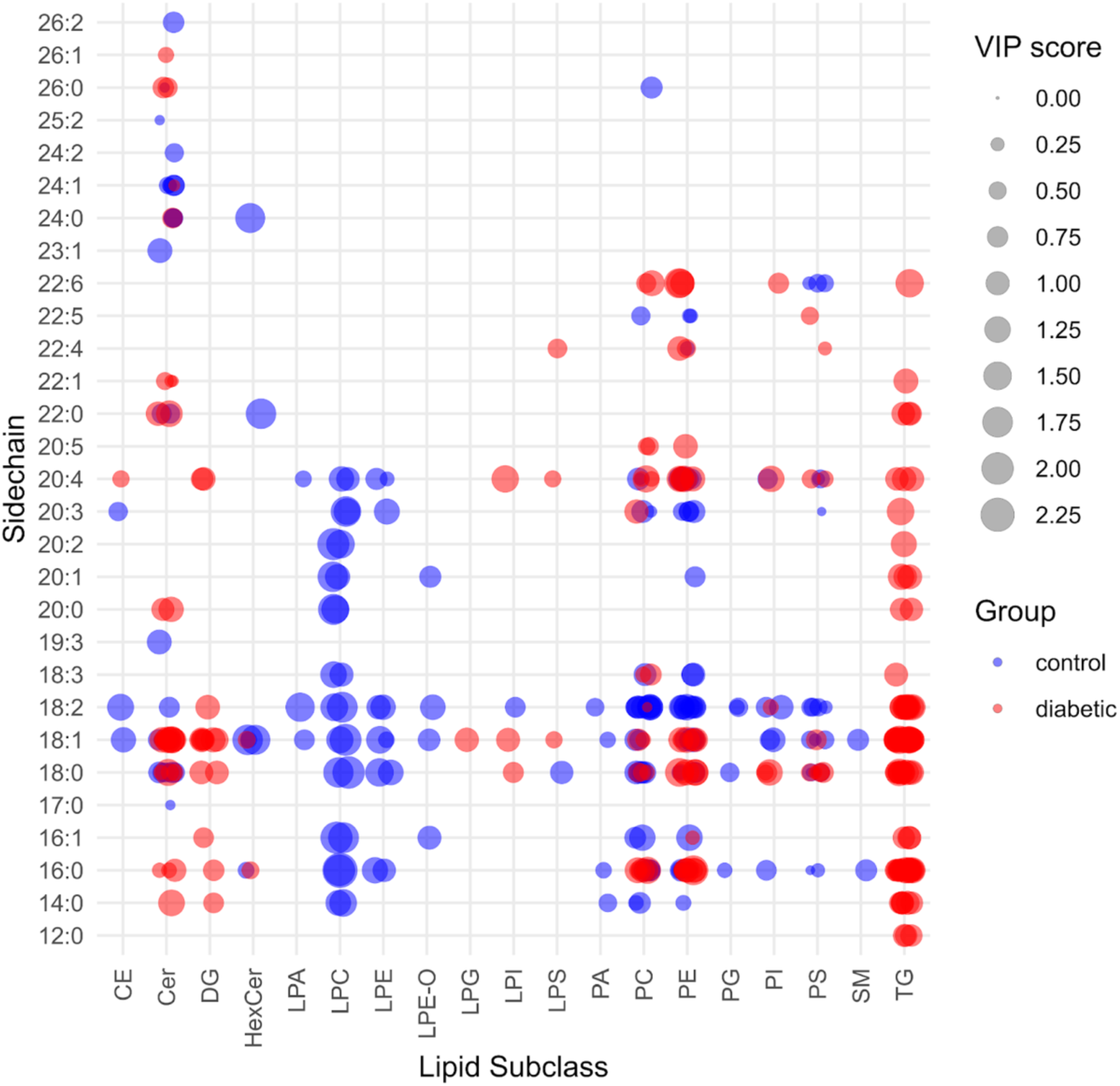
VIP scores of DBS lipids identified at the fatty acyl level based on individual lipid side chain(s) and subclass. Point size represents the magnitude of the VIP score, while point colour reflects the direction of group association based on the OPLS-DA predictive component loadings (Cohen’s d). Red indicates greater association with diabetic profiles, and blue indicates greater association with non-diabetic control profiles.

To assess how the overall degree of lipid unsaturation and total acyl carbon number influence diabetes, DBS lipids were examined at the summed species level (Figure 8). Higher degrees of unsaturation in glycerolipids such as TG and DG subclasses (e.g., TG(18:1_18:2_20:4) or TG(56:7)) were observed to skew towards higher VIP scores (consistent with the diabetic phenotype). Some shorter-chain TGs appear more neutral, but overall, TGs remain elevated in diabetic samples. Lipids with greater total acyl carbon content and saturated/mono-unsaturated side chains appear higher in diabetics, these included ceramides (Cer), ether-linked phosphatidylcholines (PC-O), and sphingomyelins (SM). Contrastingly, shorter polyunsaturated side chains for the same subclasses were elevated in non-diabetic controls. This suggests that degree of lipid unsaturation and total acyl carbons may be key discriminators between lipidomic profiles of diabetics and non-diabetic controls.

**Figure 8.**
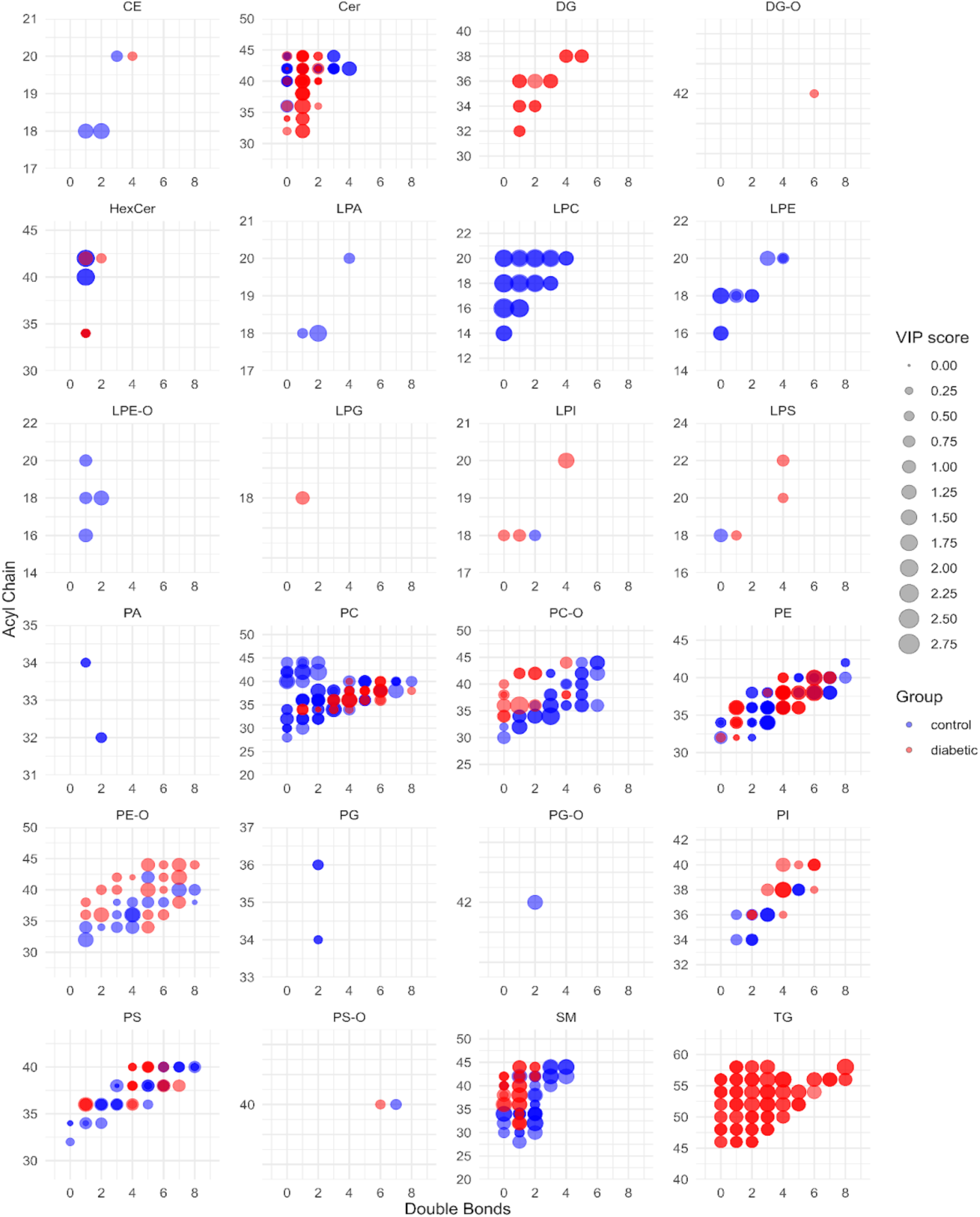
VIP scores of DBS lipids based on summed fatty acyl lipid side chains and subclass. Point size represents the magnitude of the VIP score, while point colour reflects the direction of group association based on the OPLS-DA predictive component loadings (Cohen’s d). Red indicates greater association with diabetic profiles, and blue indicates greater association with non-diabetic control profiles.

## DISCUSSION

This study evaluated the feasibility of DBS lipidomics to distinguish diabetic and non-diabetic individuals. A systematic approach was employed, including i) comprehensive semi-quantitative analysis of 23 lipid subclasses, ii) multivariate statistical modelling (OPLS-DA) to assess lipidomic differences across DBS, venous plasma, and capillary plasma samples, and iii) identification of key lipid species driving lipidomic separation, focusing on acyl chain composition and degree of unsaturation. Our findings demonstrate that DBS-derived lipid profiles align with those from traditional venous plasma, supporting the potential for decentralised, patient-centric lipidomic monitoring in diabetes research.

### Comparability of biofluids using the LC-timsTOF-MS workflow

Lipidomic profiling of self-collected DBS samples effectively differentiated diabetic and non-diabetic participants. The performance of OPLS-DA modelling across different sample matrices demonstrated the capability of self-collected DBS microsamples to preserve lipid profiles comparable to those derived from traditional venous plasma. The high R²Y (0.82) and Q²Y (0.50) values (both p = 0.05) indicate strong explanatory power and predictive ability, reinforcing the reliability of DBS in distinguishing lipidomic variations between diabetic and non-diabetic individuals. Further, key lipid species included: HexCer(18:1;O2/24:0), HexCer(18:1;O2/22:0), LPC(16:0), PC(O-34:3), TG(18:0_18:1_20:3), and TG(18:1_18:1_22:6), which were consistently identified in the top 12 VIP scores across all biological matrices (Figure 6), reinforcing the robustness of the DBS platform. These findings extend previous work using capillary plasma for lipoproteins and lipid analysis^49, 50^, by demonstrating similar diagnostic resolution from 10 µL of whole blood without plasma separation. Compared to previous studies that primarily rely on venous blood for lipidomic analyses in diabetes^51–55^, DBS offer a simplified, scalable workflow with minimal sample processing, which may offer a more accessible and patient-friendly approach to lipid monitoring for diabetes, using participant self-collection.

### Biological concordance with literature for potential therapeutic monitoring

Class-specific lipid differences in DBS samples were evident between diabetic and non-diabetic groups, particularly in glycerolipids such as TGs and DGs. Diabetic individuals exhibit an enrichment of TGs. In particular, those with longer and more unsaturated side chains, such as C22:6 and C20:3. This aligns with established links between TG accumulation, insulin resistance, and metabolic dysfunction, where impaired lipid metabolism contributes to altered energy storage and utilisation in diabetes^13, 14, 56, 57^. In contrast, LPCs were higher in non-diabetic controls compared to diabetics (Figure 7) which may indicate roles in lipid signalling. Whilst identifying the specific pathophysiological mechanisms was not within the scope nor focus of the present study, previous research has indicated both pro-inflammatory^58^ and anti-inflammatory^59^ effects of LPCs. Additionally, shorter saturated and monounsaturated fatty acid side chains, including C14:0, C16:0, C16:1, C18:0, and C18:1, were also higher in non-diabetic controls, possibly reflecting differences in lipid turnover and oxidation pathways^60–63^. Notably, Cer and SM lipids were elevated in diabetic individuals, particularly those with longer saturated and monounsaturated acyl chains, consistent with their role in promoting insulin resistance and inflammatory responses that have been previously identified^64–67^. These findings highlight lipidomic alterations captured as part of this preliminary study that successfully stratify diabetic participants and non-diabetic controls, and highlight potential mechanisms that could contribute to disease progression.

Lipid structural characteristics, specifically total acyl chain carbon content and degree of unsaturation (Figure 8), play a key role in shaping lipidomic differences between diabetic and non-diabetic individuals. In diabetes, polyunsaturated long-chain TGs, such as TG(56:7) and TG(58:8), were more abundant, reflecting their association with lipotoxicity^68^ and altered lipid turnover in insulin resistance^57^. This aligns with broader trends showing that polyunsaturated TGs and diacylglycerols (DGs) with higher total acyl chain carbon content are elevated in diabetes, indicating disrupted lipid metabolism^51^. In contrast, lipids with lower total carbon content and higher unsaturation, particularly within lyso-glycerophospholipid and sphingolipid subclasses, are more commonly observed in non-diabetic controls, potentially reflecting a protective lipidomic profile. Notably, lower levels of bioactive lysophosphatidylcholine (LPC) species have been linked to increased cardiometabolic risk^4–7, 69–72^. These findings are consistent with the underlying pathophysiology of diabetes, including impaired fasting glucose, reduced cellular glucose uptake, and insulin resistance, and are supported by prior plasma lipidomic studies that report an inverse relationship between LPC levels and obesity or type 2 diabetes^73–75^. Notably, LPC(16:0) has been shown to enhance glucose uptake independently of insulin by stimulating adipocyte glucose uptake^76^, and reductions in LPCs have been consistently documented across diabetic cohorts^4, 77–80^. Collectively, these data support a model in which lipid side chain composition contributes to metabolic phenotyping of DBS samples which may inform cardiometabolic monitoring in diabetes.

### Implications for patient-centric sampling and decentralisation

These findings have several translation implications; i) low-volume DBS sampling for monitoring of lipids in relation to diabetes complications, and ii) early identification of lipid signatures associated with the incidence and progression of diabetic complications to complement current clinical risk predictors. Preliminary findings in the present research support translation to such micro-scale DBS blood samples that can be self-collected. For this purpose, self-collected DBS offer a promising tool for real-world lipidomic monitoring, providing a minimally-invasive and accessible alternative to traditional venous blood sampling. Positive attitudes toward sample self-collection in the present study, highlight the practicality and acceptability of microsampling for lipidomics in home settings. Specifically, the perception of participants to aspects of microsampling, including preference over a monthly traditional phlebotomy draw (“strongly preferred” on average), and pain from performing a finger-prick (average <2 on a 10 point scale), demonstrate this (Figure 3). Findings by Van Uytfanghe et al.^31^, who originated the survey, similarly reported positive perceptions to microsampling of study participants, which supports the feasibility of patient-centric and decentralised sampling by non-experienced individuals. Additional data captured as part of the perceptions of microsampling survey was in agreement with existing evidence for the connection of education and diabetes^81–84^. Recently, DBS-based lipidomics has demonstrated that the majority of lipid classes remain stable under ambient and conventional fridge/freezer storage environments for up to 2 weeks^30^. Taken together with the findings from the present research, this highlights the suitability of DBS for decentralised sample collection in lipidomic research workflows. Unlike existing technologies such as CGM, which tracks short-term glucose fluctuations, DBS can enable broader lipidomic assessment. Integrating DBS-based sampling alongside personalised diabetes management strategies could enhance disease monitoring by allowing more frequent assessment of lipid alterations, tracking disease progression, and evaluating responses to interventions. Future research must focus on validating DBS-based lipidomics in larger cohorts and exploring the potential for stratification of diabetic subgroups (type 1, type 2, prediabetes, gestational diabetes, and their associated complications) based on the lipidome to refine precision diabetes care.

## CONCLUSION

This study demonstrates the feasibility of self-collected dried blood spots (DBS) for lipidomic profiling in diabetes research as a viable patient-centric alternative to traditional venous plasma sampling. Using an optimised untargeted LC-timsTOF-MS lipidomics workflow, we demonstrate the robustness of DBS samples for distinguishing lipidomic profiles between diabetic participants and non-diabetic controls. DBS samples revealed significant lipidomic differences between diabetic and non-diabetic individuals, specifically glycerolipid (e.g., TG, DG) enrichment in diabetics and lyso-glycerophospholipid (e.g., LPC) enrichment in non-diabetics. Additionally, associations between lipidomic signatures and psychosocial factors (PAID-20 and PAM-13 scores) highlight the potential for lipidomics to provide deeper insights into diabetes management and patient behaviour. Importantly, participants reported positive perceptions of DBS self-sampling, noting the easy collection method, convenience, minimal discomfort, and greater sense of autonomy compared to conventional phlebotomy. Future studies should focus on expanding cohort sizes and validating DBS lipidomic profiles across diverse populations. Regardless, the adoption of DBS for lipidomics may enhance personalised disease monitoring and improve accessibility for lipidomic health assessments, particularly in remote and resource-limited settings. Overall, our findings support the integration of DBS microsampling into longitudinal and decentralised diabetes research. This approach enables frequent, minimally invasive sampling without the logistical barriers of venous phlebotomy, facilitating future studies on lipid biomarkers not only in diabetes risk assessment and disease progression, but also in the broader context of cardiometabolic health.

## Notes

### Competing Interest Statement

The authors have declared no competing interest.

