## Supporting Information for "Dried Blood Spot Lipidomics in Diabetes"

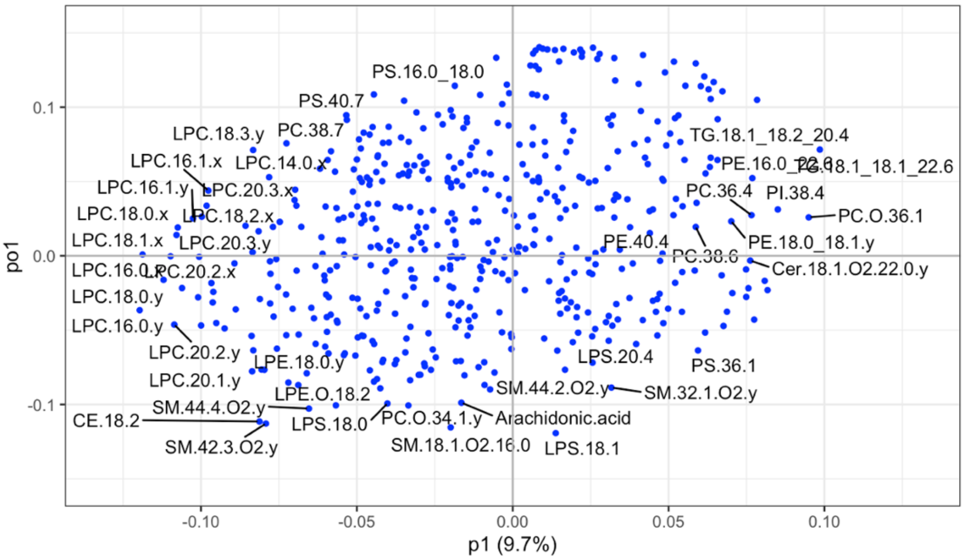


**Figure S.1. Loadings plot of the OPLS-DA model between non-diabetic controls and diabetic groups for self-collected DBS samples.**

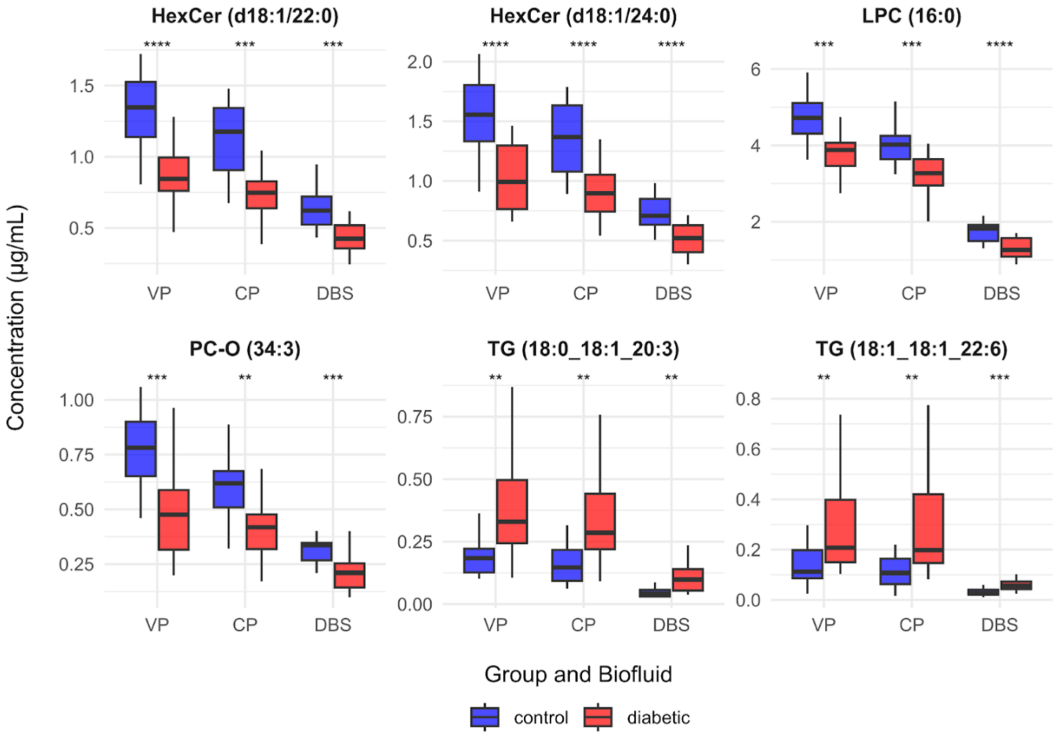


**Figure S.2. Boxplots of lipids from the OPLS-DA analysis that had consistent top VIP-scores across all biological samples.** Non-diabetic controls (blue) and diabetic groups (red). Biological sample types: venous plasma (VP), capillary plasma (CP), and dried blood spot (DBS).

**Table S.1. PAID-20 results for diabetics**

| Item | Description | Avg [+/-SD] |
|---|---|---|
| PAID_1 | Not having clear and concrete goals for your diabetes care? | 1.53 ± 0.94 |
| PAID_2 | Feeling discouraged with your diabetes treatment plan? | 1.29 ± 1.1 |
| PAID_3 | Feeling scared when you think about living with diabetes? | 1.18 ± 1.13 |
| PAID_4 | Uncomfortable social situations related to your diabetes care (e.g. people telling you what to eat)? | 1.18 ± 1.19 |
| PAID_5 | Feelings of deprivation regarding food and meals? | 1.06 ± 0.9 |
| PAID_6 | Feeling depressed when you think about living with diabetes? | 0.94 ± 0.9 |
| PAID_7 | Not knowing if your mood or feelings are related to your diabetes? | 1.06 ± 0.83 |
| PAID_8 | Feeling overwhelmed by your diabetes? | 1 ± 0.94 |
| PAID_9 | Worrying about low blood glucose reactions? | 1.47 ± 0.8 |
| PAID_10 | Feeling angry when you think about living with diabetes? | 0.76 ± 0.75 |
| PAID_11 | Feeling constantly concerned about food and eating? | 1.29 ± 0.69 |
| PAID_12 | Worrying about the future and the possibility of serious complications? | 1.41 ± 0.87 |
| PAID_13 | Feelings of guilt or anxiety when you get off track with your diabetes management? | 1.53 ± 1.01 |
| PAID_14 | Not accepting your diabetes? | 0.76 ± 1.03 |
| PAID_15 | Feeling unsatisfied with your diabetes physician? | 0.71 ± 1.16 |
| PAID_16 | Feeling that diabetes is taking up too much of your mental and physical energy every day? | 1.35 ± 1.22 |
| PAID_17 | Feeling alone with your diabetes? | 0.53 ± 0.8 |
| PAID_18 | Feeling that your friends and family are not supportive of your diabetes management efforts? | 0.76 ± 1.03 |
| PAID_19 | Coping with complications of diabetes? | 1.18 ± 0.95 |
| PAID_20 | Feeling burned out by the constant effort needed to manage diabetes? | 1.41 ± 1.18 |
| PAID_Raw | Raw score /80 | 22.41 ± 13.24 |
| PAID_Score | Raw score * 1.25 | 28.01 ± 16.55 |
| PAID_level | Level | 1.94 ± 0.66 |
| **PAID subscales*** | | |
| Diabetes-related emotional distress (PAID items 3, 6, 7, 8 , 9, 10, 12, 13, 14, 16, 19, 20) | | 1.17 ± 0.69 |
| Treatment-related problems (PAID items 1, 2, 15) | | 1.18 ± 0.85 |
| Food-related problems (PAID items 4, 5, 11) | | 1.18 ± 0.7 |
| Social support-related problems (PAID items 17, 18) | | 0.65 ± 0.81 |

Note. Likert scale ranges 0 (not a problem) – 4 (serious problem). ***PAID subscales** as proposed by *F J Snoek, F Pouwer, G W Welch, W H Polonsky; Diabetes-related emotional distress in Dutch and U.S. diabetic patients: cross-cultural validity of the problem areas in diabetes scale.. Diabetes Care 1 September 2000; 23 (9): 1305–1309. https://doi.org/10.2337/diacare.23.9.1305*
